# Inhibition of *Fusarium oxysporum* growth in banana by silver nanoparticles: *in vitro* and *in vivo* assays

**DOI:** 10.1101/2024.07.22.604513

**Authors:** Natalia Mendoza, Paola Yánez, Freddy Magdama, Ricardo Pacheco, Joel Vielma, María Eulalia Vanegas, Nina Bogdanchikova, Alexey Pestryakov, Pablo Chong

**Affiliations:** ESPOL Polytechnic University, ESPOL, Centro de Investigaciones Biotecnológicas del Ecuador, Campus Gustavo Galindo, Km 30.5 Vía Perimetral, Guayaquil, 090902, Ecuador; ESPOL Polytechnic University, ESPOL, Facultad de Ciencias Naturales y Matemáticas, Departamento de Ciencias Químicas y Ambientales, Campus Gustavo Galindo, Km 30.5 Vía Perimetral, Guayaquil, 090902, Ecuador; Center for Environmental Studies (N@NO-CEA Group), Department of Applied Chemistry and Production Systems, Faculty of Chemical Sciences, University of Cuenca, 010203 Cuenca, Ecuador; Centro de Nanociencias y Nanotecnología, Universidad Nacional Autónoma de México, Km 107 Carretera Tijuana-Ensenada, 22860, Ensenada, BC, México; Research School of Chemistry and Applied Biomedical Sciences, Tomsk Polytechnic University, 634050 Tomsk, Russia

**Keywords:** Silver nanoparticles, *Musa*, *Fusarium oxysporum* f. sp. *cubense*, Pathogen control

## Abstract

Fusarium wilt is a devastating disease that affects banana crops worldwide. In Ecuador, bananas are one of the most important commodities and staple food. Nanoparticles are emerging as innovative solutions to control fungal diseases in plant protection. In this study, *in vitro* and *in vivo* assays were carried out to validate *Fusarium oxysporum* growth and disease inhibition. 96-well plates experiments were used to calculate the IC_50_ of three different silver nanoparticle formulations (Argovit-1220, Argovit-1221, and Argovit-C) against four Ecuadorian *Fusarium* strains (EC15-E-GM1, EC19-LR-GM3, EC35-G-GM6, EC40-M-GM2). More than 95% inhibition rate was obtained at 25 mg L^-1^ concentration. Fusarium wilt *in vivo* assay was carried out with Gros Michel plants, where better control was obtained by applying silver nanoparticles to the roots, reducing disease development by an average of 68%. This study shows that silver nanoparticles have a high antifungal potential for controlling the Fusarium wilt of bananas. To the best of our knowledge, this is the first study to test the potential of silver nanoparticles against *Fusarium oxysporum* in planta under greenhouse conditions.

## Introduction

Bananas are one of the most important crops worldwide. The fruit is an important staple food and an essential source of nutrition, rich in potassium, vitamins, and dietary fiber[1]. It is also a primary determinant of economic growth, employment, and cultural heritage in many countries. For Ecuador, in particular, bananas are the main exported agricultural product and consumption commodity[2]. The most important emerging threat to the banana industry is the Fusarium wilt caused by the soil-borne pathogen *Fusarium oxysporum* f. sp. *cubense* (Foc)[3]. Foc affects the plant vascular system, causing wilting and yellowing that eventually leads to the plant death[4]. In the 1960s Foc race 1 cost significant losses to the banana industry and a profound shift in the way bananas were cultivated[5]. The main change was the replacement of the Gros Michael cultivar with varieties belonging to the Cavendish subgroup resistant to Foc race 1. Currently, the banana industry is threatened again by a new race of Foc, colloquially known as tropical race 4 (TR4), and capable of causing disease in more than 60% of all varieties of banana in the world, including Cavendish cultivars[5]. The monoculture of bananas, its genetic uniformity, the non-availability of chemical control, and the lack of proper phytosanitary national policies guarantee the failure of disease management, biosafety, and quarantine procedures[3]. Although a new recombinant banana *Fusarium*-resistant cultivar has been liberated in Australia[6], it is not broadly available, and some countries like Ecuador have imposed legal restrictions on the adoption of certain technologies, including recombinant pathogen-resistant banana varieties[7,8], leaving no other effective control measures for this disease. These reasons emphasize the urgency to develop new solutions for *Fusarium* control in banana plantations.

New alternative technologies, such as nanotechnologies are rising for multiple biological applications. Based on their antimicrobial properties nanoparticles (NPs) have been widely used in many agricultural applications[9]. The advantages of the NPs for plant pathogens control are their low-dose effectiveness[9]. Silver (Ag) and its derivatives are acknowledged for their antimicrobial properties[10,11]. Silver nanoparticle (AgNP) modes of action include the disruption of the microbe cell membrane potential[10] or the inhibition of processes like DNA or RNA synthesis, ultimately leading to cell death[12]. AgNPs have been proven to be effective against *Botrytis cinerea*, *Alternaria alternata*, *Monilinia fruticola*, *Colletotrichun gloeosporioides*, and *Fusarium oxysporum* f. sp*. Radices-Lycopercisi*, *Fusarium solani* and *Verticillium dahlia* on both *in vitro* and *in vivo* tests[11]. In particular, AgNPs have been shown to be particularly lethal at low doses to different *Fusarium* species[11]. Mahdizadeh et al. (2015) studied AgNPs against many pathogenic and beneficial fungi, showing a differential dose effect for each species, with *Trichoderma harzianum* the least affected. In this case, Mahdizadeh et al. (2015) argue that the beneficial fungi *T. harzianum* could be protected using a proper dose that will control the pathogens and simultaneously have a milder effect on the beneficial fungi growth[10].

In the present study, we assessed the efficacy of AgNPs to control Fusarium wilt on bananas. Our study shows *in vitro* control over the pathogen growth at AgNP doses as low as 25 mg L^-1^ and a substantial reduction of the disease severity in a greenhouse experiment with just one AgNP application at 100 mg L^-1^.

## Materials and Methods

### Fungal Strains

*Fusarium oxysporum* f. sp. *cubense* (Foc) race 1 strains EC15-E-GM1, EC19-LR-GM3, EC35-G-GM6, and EC40-M-GM2 were provided by the collections of the Biotechnology Research Centre of Ecuador (CIBE) from ESPOL Polytechnic University.

### Culture conditions for in vitro assays

Strains were cultivated from the stock water suspensions on potato dextrose agar (PDA) and incubated at 28°C for 3 days. Cultures grown on PDA were covered with 5 - 9 mL of Tween 20 (0.05%), and the inoculum suspension was harvested by a gentle but exhaustive surface scraping with a sterile loop. Then, the obtained solution was transferred to a 50 mL conical tube. Aliquots of the solution obtained at 1:10 and 1:100 were prepared and quantified under the microscope cell-counting with a Neubauer chamber.

### AgNPs synthesis and formulations

All three AgNP samples, codes Argovit-1220, Argovit-1221, and Argovit-C, were lots based on Argovit^TM^ AgNPs (Research and Production Center “Vector-Vita” Ltd, Novosibirsk, Russia). Argovit^TM^ is a water suspension of AgNPs. The metallic content of Ag is 12 mg mL^-1^ (1.2 wt. %) with 188 mg mL^-1^ (18.8 wt. %) of the stabilizer with a final concentration of AgNPs 200 mg mL^-1^ (20 wt. %) suspension. All stocks and aliquots were stored in the dark at 4°C. The characterization of the AgNP samples is shown in Table 1.

**Table 1.** The origin of the stabilizers and physicochemical properties of AgNP samples studied at the present work.

| Sample | AgNP formulation | Stabilizer |  | Content, wt. % |  | Hydrodynamic diameter, nm | Z potential, mV | AgNP characteristics according to HRTEM data** |  | Peak positions in UV-Vis, nm |
| --- | --- | --- | --- | --- | --- | --- | --- | --- | --- | --- |
|  |  | Manufacturer | Type | Ag | Stabilizer |  |  | The average diameter, nm | Morphology |  |
| Argovit 1220 | Argovit<sup>TM</sup> | Boai NKY Pharmaceuticals Ltd., (Jiaozuo, China) | PVP* with glucose additive | 1.2 | 18.8 | 141.0 | +12.4 | Bimodal distribution with maxima at 8 nm and 80 nm | 8 nm spheroidal, 80 nm pyramidal | 412 and 437 |
| Argovit 1221 | Bioargovit<sup>TM</sup> | Russian manufacturers producing in accordance with interstate standard | Edible gelatin/hydrolyzed protein | 1.2 | 18.8 | 114.1 | -7.15 | 8.5±3.3 | Spheroidal | 420 |
| Argovit C | Argovit-C<sup>TM</sup> | Boai NKY Pharmaceuticals Ltd., and Russian manufacturers fabricating HP in accordance with interstate standard | Mixture of PVP* (1/3) and hydrolyzed protein (2/3) | 1.2 | 18.8 | 142.6 | +9.15 | 14.95±10.1 | Spheroidal | 424 and 461 |

\* PVP is polyvinylpyrrolidone
\*\*HRTEM data was taken from the previous publication of our group[13]

### Fungal in vitro growth inhibition

A 96 microwell plate was used to perform growth inhibition bioassays of Foc race 1 with AgNPs. The antifungal properties of AgNPs were reproduced on three AgNP samples and at four strains of Foc. The total volume of each well was 100 µL, which was comprised of 75 µL of Mueller Hinton Broth (Titan Biotech Ltd., Rajasthan, India) medium, 15 µL of AgNPs, and 10 µl of each fungal strain. The medium was dispensed, and the nanoparticles were dosed at different concentrations; 0, 0.8, 1.6, 3.1, 6.3, 12.5, 25, 50, and 100 mg L^-1^ calculated for metallic Ag with stabilizer. Each strain was added at a concentration of 1x10^4^ spores per millilitre. The plate was incubated for 3 days at 28°C to ensure optimal fungus growth. The compound alamarBlue^TM^ HS Cell Viability Reagent (Thermo Fisher Scientific Inc., Waltham, Massachusetts, USA) was added aseptically at 10% of the total well volume, to measure the cell viability for each treatment and determine its half-maximal inhibitory concentration (IC_50_). The study was conducted in triplicate and with three biological replicates.

The percentage of inhibition was evaluated by measuring cell viability with the alamarBlue^TM^ compound. In this reaction, the molecule resazurin (non-fluorescent blue colour) is reduced to resorufin (fluorescent pink colour)[14]. Then the percentage of inhibition was determined according to the manufacturer’s protocol[14]. IC_50_ was calculated to determine the AgNP antifungal potential, and the GraphPad Prism program version 10.0.2 (Home – GraphPad Software, Boston, Massachusetts USA, www.graphpad.com) was used for this purpose.

### Preparation of the inoculum for in vivo assays

In a 1L flask, a solution with 700 mL of filtered water and 5.6 g of mung bean was placed and autoclaved for 15 min at 121 °C with aluminium foil. Once ready, the solution was kept at room temperature until its temperature was reduced to approximately 40 °C. Five mycelium pieces (0.5 x 0.5 cm) of Foc race 1 were placed in the solution and the flask nozzle was sealed with cotton plugs. The solution with the pieces of mycelium was incubated for 8 days at 120 RPM and 25 ±2 °C. Once the incubation period was over, the solution was filtered using a vacuum pump and a double layer of sieving gauze. The filtrated solution with spores was stored at 4 °C until its later use. A diluted spore solution at 1:100 was used three times to estimate spore concentration with a Neubauer cell counting chamber (Boeco, Hamburg, Germany). The final solution was adjusted to 1x10^4^ spores per millilitre.

### Greenhouse bioassay conditions

Under greenhouse conditions, 84 three-month-old Gros Michel banana plants were set on a substrate composed of compost, sand, and rice husks in a 6:2:2 ratio, for the *in vivo* bioassays. The experiments were conducted with five replicates per treatment. Two methods of infection were employed for the experiments: drenching and root dipping. For infection by drenching, 100 mL of a concentrated inoculum solution at 2 x10^5^ spores per millilitre was poured into the substrate. Expanded polystyrene plates were placed under the pots to keep the spore solution in contact with the substrate. For the infection by root dipping, the roots were separated from the substrate to be soaked in the inoculum solution for 30 - 40 min. Once this period ended, the plants with the infected roots were planted again in the fertilized substrate.

After 72h of infection, the plants were watered and treated with AgNP samples (Argovit-1220, Argovit-1221, and Argovit-C) at 100 mg L^-1^. The AgNPs were applied in separated experiments, one to the leaves and one to the roots. The plants were kept under evaluation for 40 days.

### Evaluation of Antifungal Activity

The software ImageJ was used to quantify the corm-affected area by Foc race1 following the methodology of Pride et al. (2020)[15]. The selection of the corm-healthy and necrotic tissue area was obtained by adjusting and changing the colour threshold parameters (hue, saturation, and brightness) to measure each area. The following formula described by Islam and Islam (2015)[16] was used to calculate the percentage of the infected area in each plant:

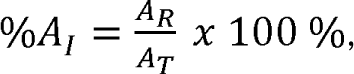

where A_R_ is the area of the affected region of the corm, and A_T_ is the total area of the corm.

### Statistical Analysis

The data in this study was analyzed based on a comparison with the treatments and their respective control data, using ANOVA. Statistical significance difference was considered when p values were less than 0.05. All the tests were performed with the software InfoStat (version 2008, F.C.A-U.N.C, Argentina). After the analysis, the data were log_2_-transformed to show a better visual scale representation in some figures.

## Results

### In vitro bioassays of Fusarium interaction with AgNPs

The results obtained from inhibition tests performed *in vitro* showed that AgNPs have an antifungal effect against Foc race 1 strains. Fig 1 presents the mycelial growth inhibition percentage obtained in the experiment for each treatment tested. All three AgNP samples show antifungal activity against all the strains at different AgNP concentrations. Most strains reached approximately 50% fungal growth inhibition at concentrations around 3.1 mg L^-1^ and 6.3 mg L^-1^. More than 90% inhibition was achieved at concentrations from 25 mg L^-1^ to 50 mg L^-1^. The studied strains responded differently to the AgNPs. The strain Ec35-G-GM6 showed the lowest sensitivity toward the AgNP samples (S1 Table).

**Fig 1.**
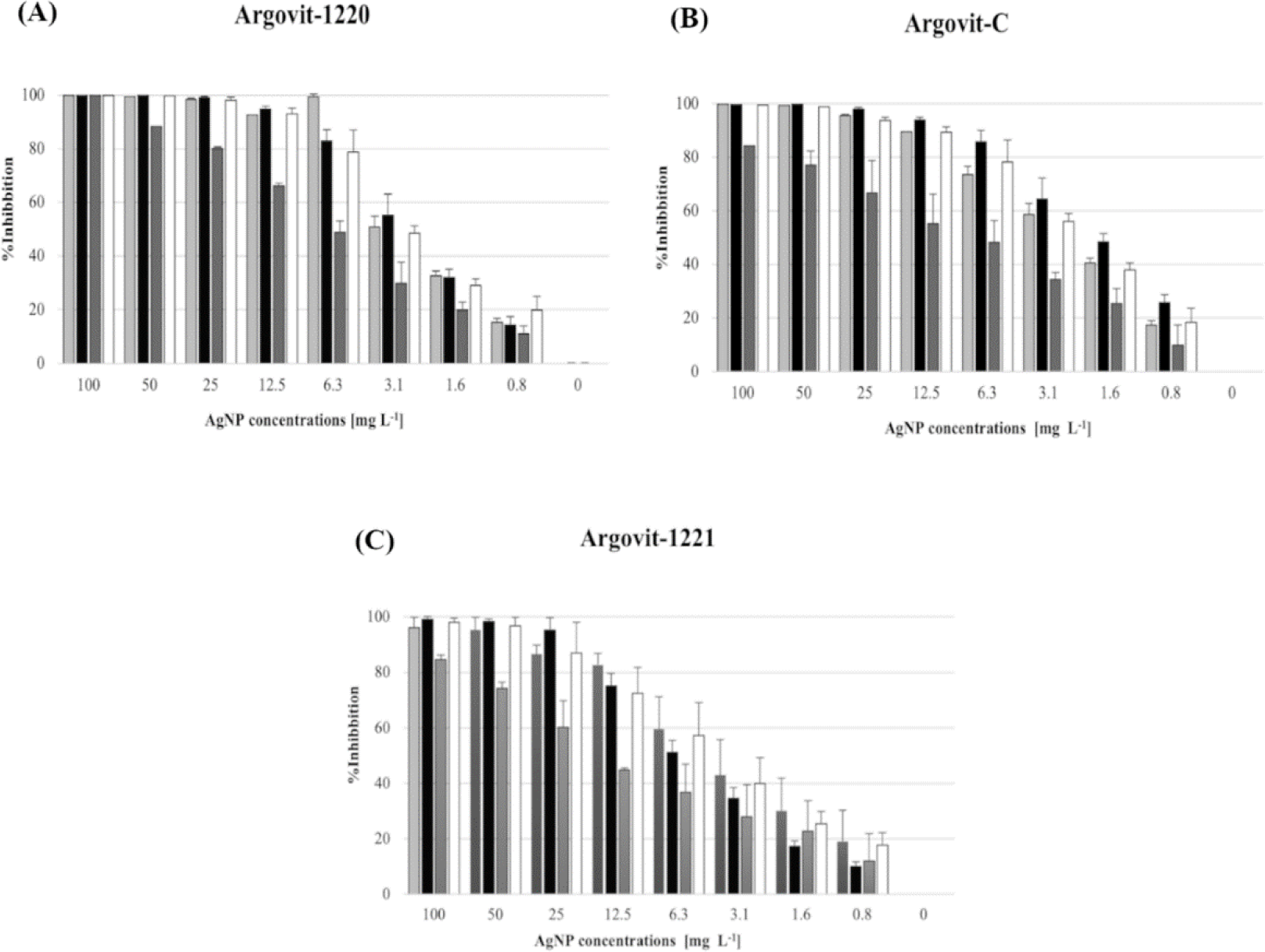
Results of the percentages of growth inhibition of Foc race 1 (4 strains) at different AgNP concentrations for three AgNP samples. Bar diagrams shows the effect of the AgNP samples on *Fusarium* inhibition growth, where (A) displays the Argovit-1220, (B) displays the Argovit-C, and, (C) displays the Argovit-1221 results.

Fig 2 shows the calculated IC_50_ values from the strainś responses. The IC_50_ values from the interaction of Argovit-1220 with EC15-E-GM1, EC19-LR-GM3, and EC40-M-GM2 strains were in the range of 3.00 mg L^-1^ – 3.75 mg L^-1^. Remarkably, for the EC35-G-GM6 strain, the value was almost double (7.39 mg L^-1^). For the Argovit-1221 sample, the IC_50_ values with EC15-E-GM1, EC19-LR-GM3, and EC40-M-GM2 were from 4.49 mg L^-1^ – 6.66 mg L^-1^, but for the EC35-G-GM6 strain, the IC_50_ was almost triple (23.26 mg L^-1^). For Argovit-C, the IC_50_ for EC15-E-GM1, EC19-LR-GM3, and EC40-M-GM2 varied from 1.35 mg L^-1^ to 2.12 mg L^-1^, again for the EC35-G-GM6 strain required almost double concentration (4.23 mg L^-1^).

**Fig 2.**
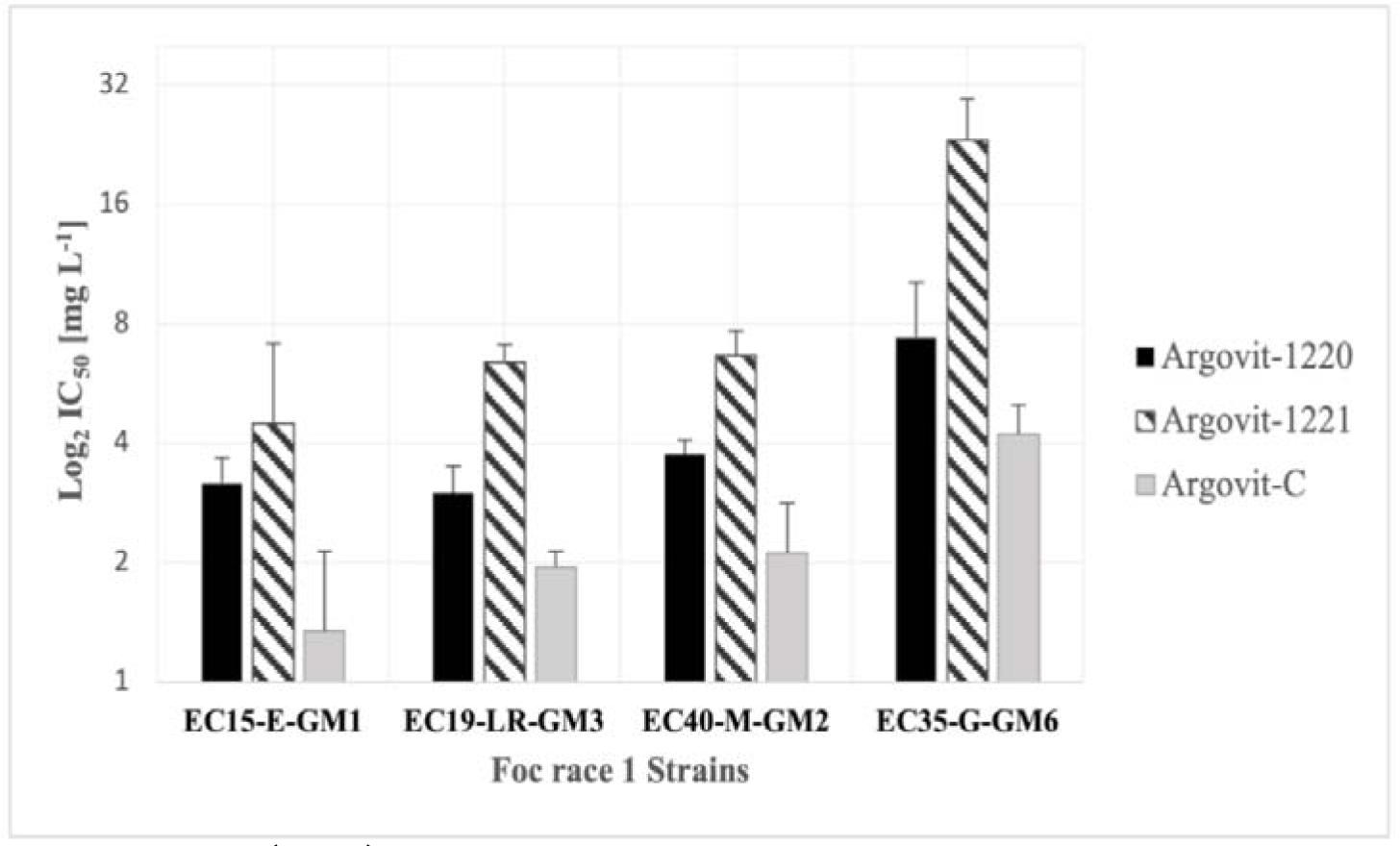
Means IC_50_ (Log_2_) values of the AgNP samples. Values of the half inhibitory concentrations for the Argovit-1220, Argovit-1221, and Argovit-C interacting with Foc race 1 strain: EC15-E-GM1, EC19-LR-GM3, EC35-G-GM6, and EC40-M-GM2.

It is observed that each strain has a distinct sensitivity to the treatments. Fig 3 shows that EC35-G-GM6 demonstrated the highest average concentration of IC_50_ (11.63 ± 9.47 mg L^-1^), indicating a significant difference between the other strains’ responses: EC15-E-GM1 (3.15 ± 1.78 mg L^-1^), EC19-LR-GM3 (3.79 ± 2.07 mg L^-1^), and EC40-M-GM2 (4.18 ± 2.10 mg L^-1^). However, strains EC15-E-GM1, EC19-LR-GM3, and EC40-M-GM2 showed not statistically different between their IC_50_ values.

**Fig 3.**
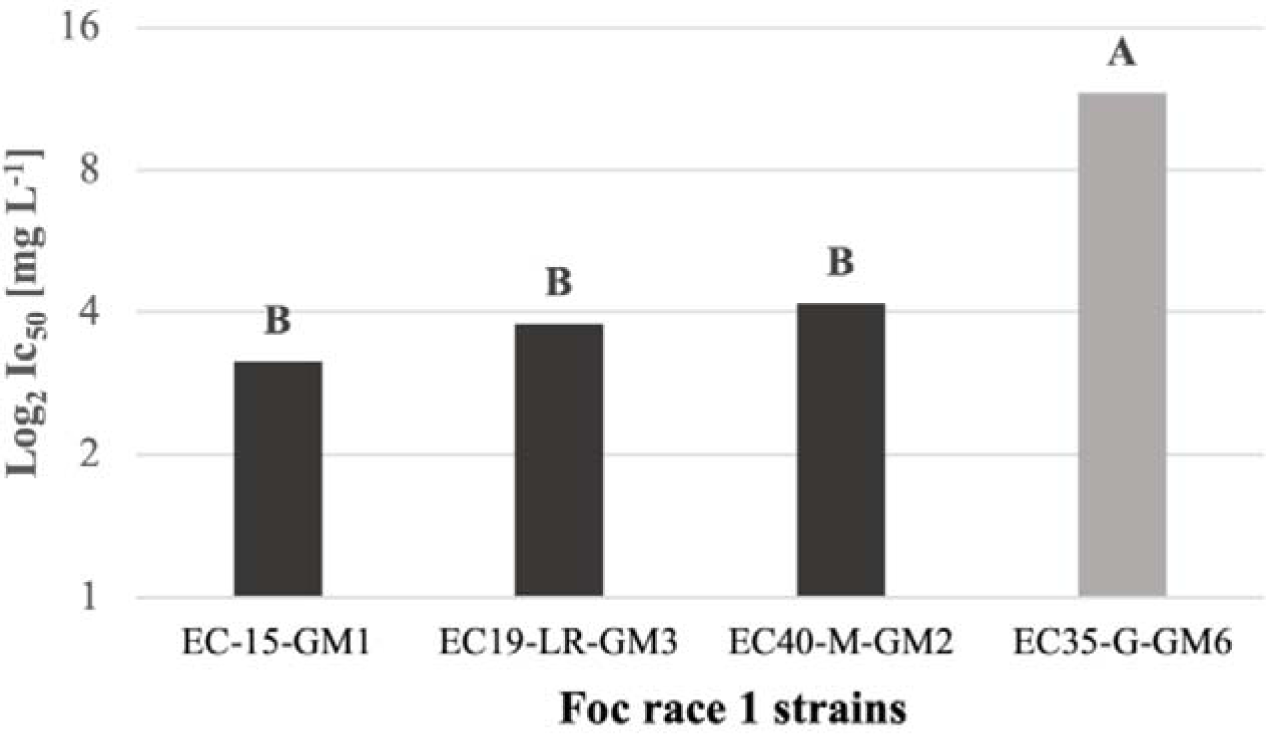
IC_50_ (Log_2_) combined means values of the different Foc race 1 strains. The letters A and B represent the statistically significant difference between them according to the ANOVA test (p > 0.05).

In Fig 4, on average, Argovit-1221 appeared as the least effective AgNP sample as it had a higher IC_50_ value (10.21 ± 8.44 mg L^-1^) than the other samples, showing a statistical difference with Argovit-1220 and Argovit-C values. However, there is no statistical difference between Argovit-1220 (4.33 ± 2.26 mg L^-1^) and Argovit-C (2.52 ± 1.13 mg L^-1^).

**Fig 4.**
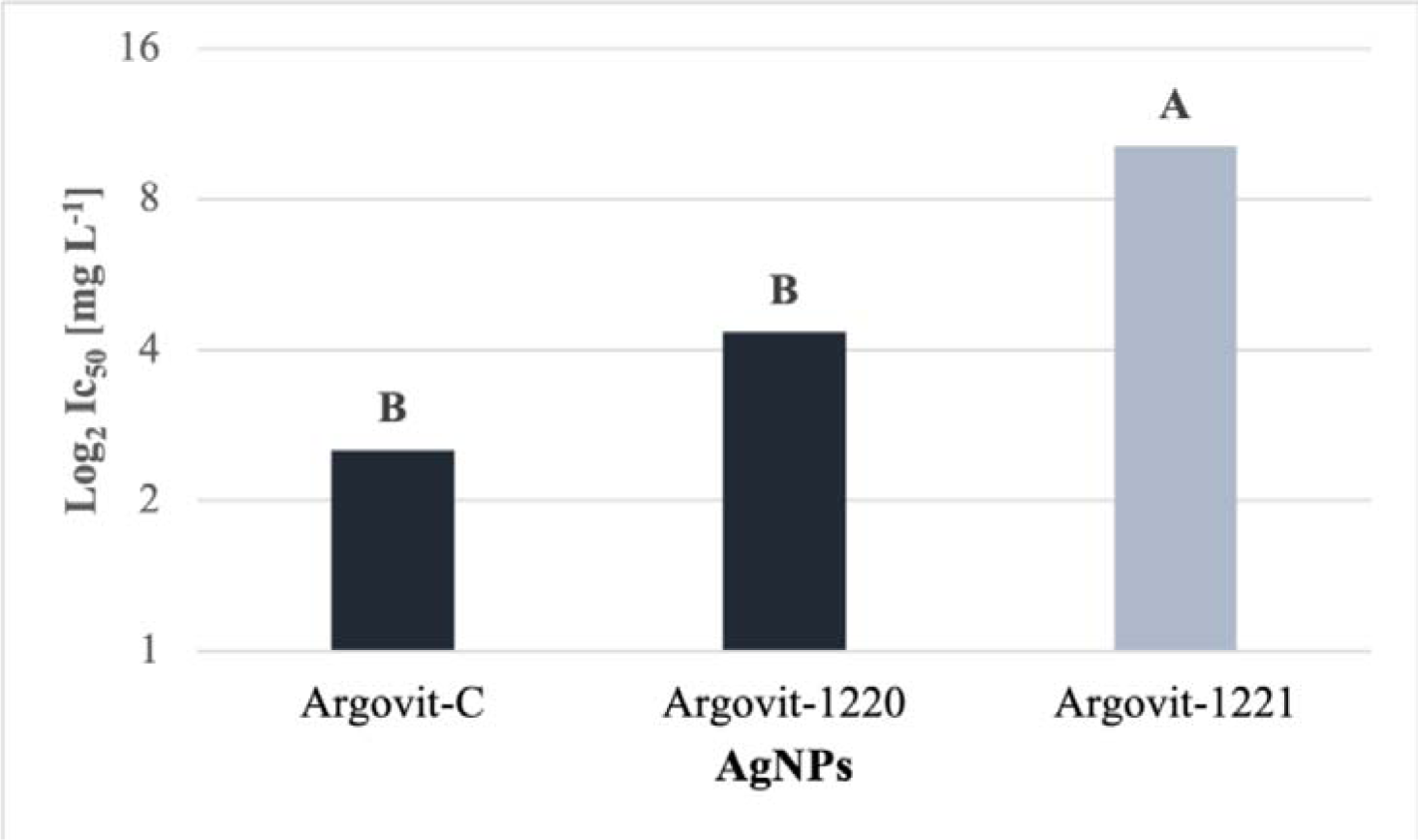
Comparison between the combined average IC_50_ (Log_2_) values of the AgNP samples for all Foc race 1 strains. The letters (A, B) mean the significant difference between them according to the ANOVA test (p > 0.05).

### In vivo bioassays of Fusarium interaction with AgNPs

The percentage of disease inhibition was determined considering the ratio of the infected area and the total area of the corm. Table 2 presents the results obtained with the radicular and foliar application of AgNPs for the two Foc infection methods (root dipping and drenching). The three AgNP samples showed good control for both modes of application. The range of inhibition obtained by the Argovit-1220 sample was from 67% to 86%. Likewise, the Argovit-C inhibition range was from 55% to 84%. The Argovit-1221 range of inhibition obtained was from 47% to 82%. Overall the Argovit-1220 treatment showed a better control than the other AgNPs-treated groups with its inhibition average of 76.73% ± 9.82. The Argovit-C sample exhibited a similar control to Argvoti-1220 with an inhibition average of 75.21% ± 13.88. The Argovit-1221 was the least effective in controlling the pathogen with an inhibition average of 67.38% ± 17.09.

**Table 2.**
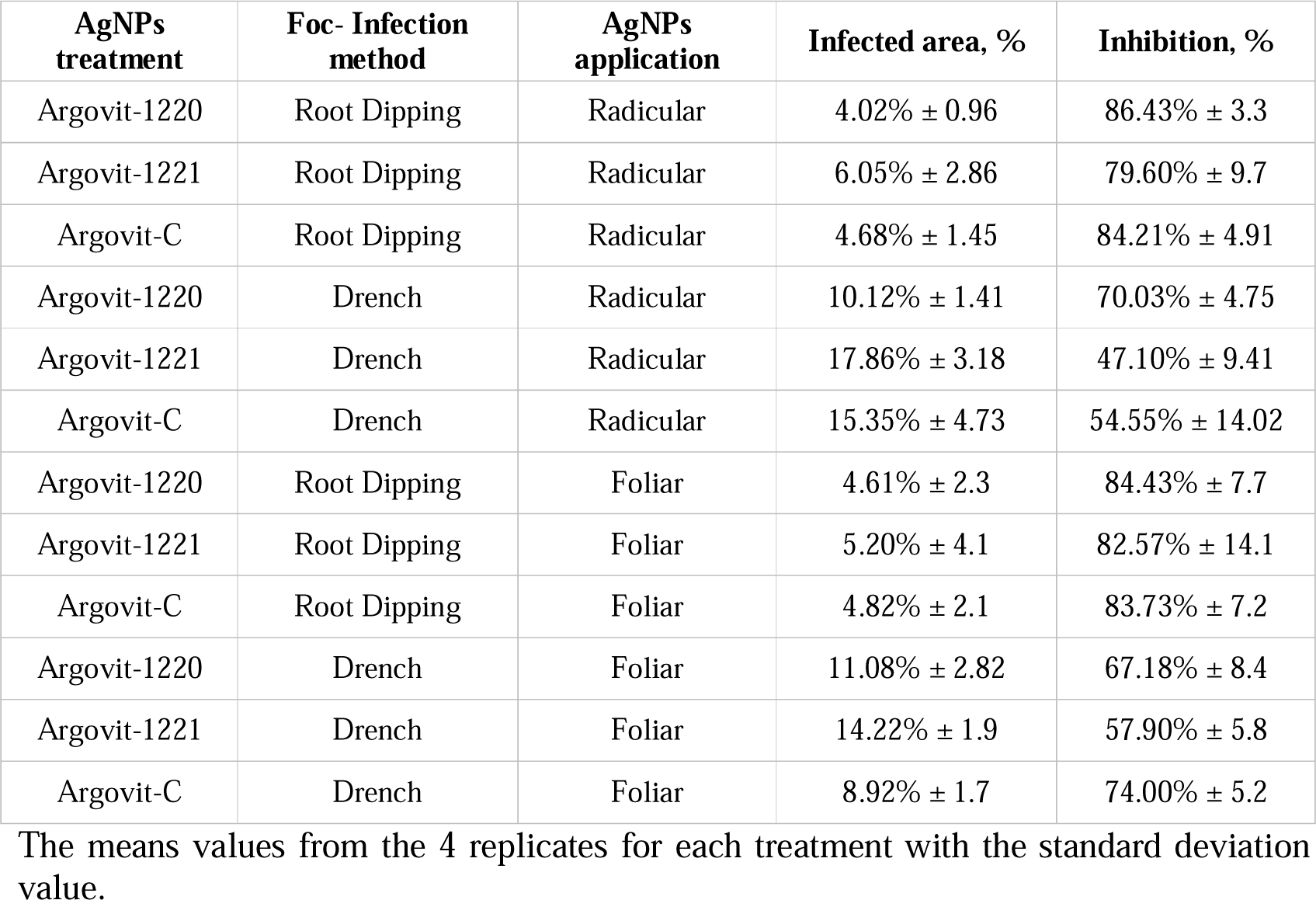
Results of Foc race 1 interaction with AgNPs in in vivo bioassay.

In relation to Table 2, Figs 5 and 6 provide a visual representation of the corm condition, with a comparison between the AgNPs treatment and their respective controls (positive and negative). The positive control is defined as the corm that has been infected with Foc race 1 but has not undergone any AgNP treatment. The negative control is the corm without the inoculum but with the application of the AgNPs excipient at the same concentration (100 mg L^-1^). The photos in Fig 5 show the results of the foliar application of AgNPs, and Fig 6 shows the results of the radicular application. The photographs exhibited that the Fusarium wilt can be controlled and held back by the three samples of silver nanoparticles (Argovit-1220, Argovit-1221, and Argovit-C) because the marked difference between the treatments and the positive controls are evident.

**Fig 5.**
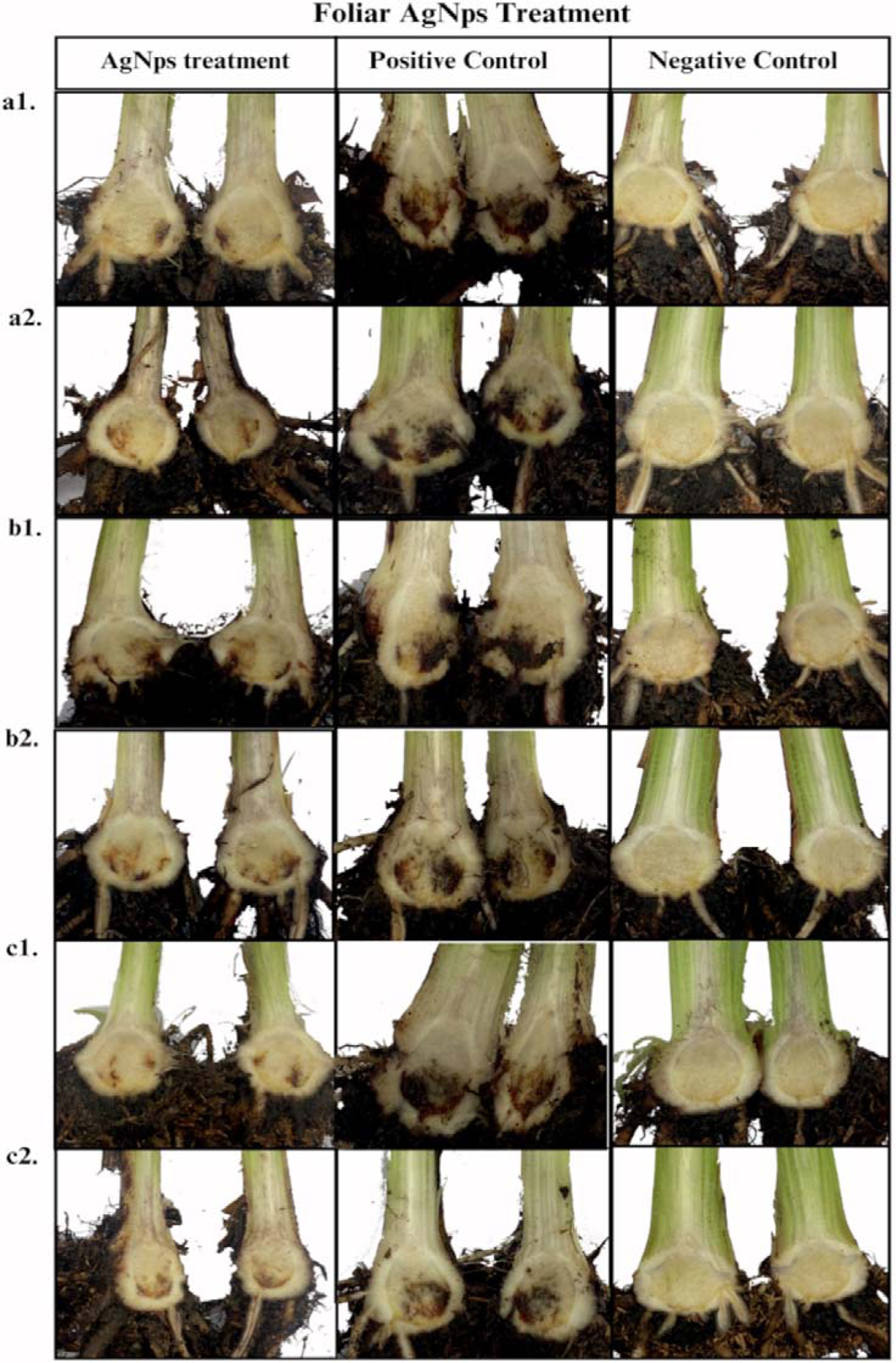
Pictures of the control of Fusarium wilt in Gros Michel plants with foliar AgNPs application. a1. Argovit-1220 with root dipping infection, a2. Argovit-1220 with drench infection, b1. Argovit-1221 with root dipping infection, b2. Argovit-1221 with drench infection, c1. Argovit-C with root dipping infection, and c2. Argovit-C with drench infection. Positive control (no AgNP application to infected plants) and negative control (not infected plants without AgNP application).

**Fig 6.**
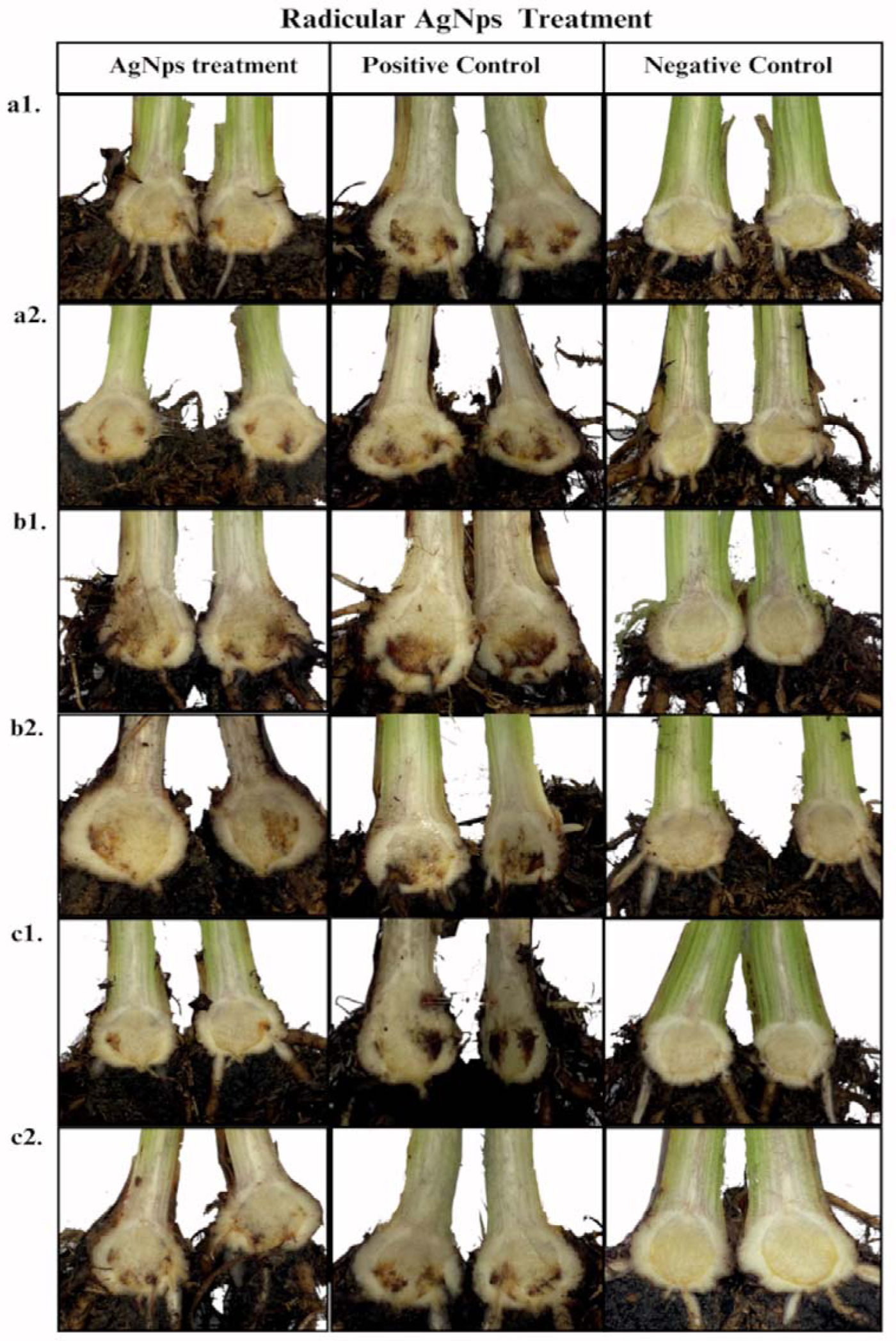
Control of Fusarium wilt in Gros Michel plants with AgNps radicular (drench) application. a1. Argovit-1220 with root dipping infection. a2. Argovit-1220 with drench infection. b1. Argovit-1221 with root dipping infection. b2. Argovit-1221 with drench infection. c1. Argovit-C with root dipping infection and c2. Argovit-C with drench infection. Positive control (no AgNP application to infected plants) and negative control (not infected plants without AgNP application).

## Discussion

### Antifungal activity in vitro

This study evaluated the antifungal potential of AgNPs against various strains of Foc race 1, a plant pathogen causing Fusarium wilt, a devastating disease in banana crops worldwide. The bioassay results demonstrated the promising antifungal activity of studied AgNP samples (Argovit-1220, Argovit-1221, and Argovit-C) against Foc race 1 strains, as shown by the low IC_50_ values obtained. The effectiveness of AgNPs as antimicrobial agents has been extensively reported in the literature[17]. AgNPs possess unique physicochemical properties that contribute to their antimicrobial activity. These include a high surface area-to-volume ratio and the ability to generate reactive oxygen species, which can lead to oxidative stress and subsequent cell death in microorganisms[18]. The study conducted by Terzioğlu et al.[19] revealed that AgNP mechanisms of action may exert their antifungal effects by inactivating certain vital enzymes, which leads to an increase in reactive oxygen species (ROS) and cellular oxidative stress, and a reduction in mitochondrial membrane potential, ultimately resulting in cell death. Another mechanism of AgNP action reported is the AgNP diffusing into the fungal cells and disrupting the microbe cell membrane[10].

Previous studies on *Fusarium oxysporum* f. sp. *radicis-lycopersici* showed that AgNPs controls Foc at a concentration of 25 mg L^-1^ and above[20]. Analysis of the IC_50_ values obtained in our experiments revealed notable differences in susceptibility to AgNPs among the tested Foc race 1 strains. Overall, lower IC_50_ values indicated higher potency, showing that lower concentrations of AgNPs were required to inhibit fungal growth by 50%. The nanoparticles that appear to have the most promising antifungal potential are the Argovit-1220 and Argovit-C samples, as they require lower concentrations to achieve effective inhibition (12.5 mg L^-1^ - 25 mg L^-1^). However, Argovit-1221 demonstrates that it requires higher concentrations (50 mg L^-1^ – 100 mg L^-1^), as evidenced by Table S1.

Interestingly, strain EC-35 exhibited the highest IC_50_ values across all three tested samples of AgNPs, indicating the lowest susceptibility to the AgNPs compared to the other strains tested. The variation in susceptibility to AgNPs among different Foc strains underscores the importance of considering strain-specific responses when evaluating antifungal agents. Some strains may exhibit reduced susceptibility to antifungal agents due to mutations, innate defence, or stress mechanisms. The study performed by Terzioğlu et al.[19] revealed that fungi respond to heavy metals, including silver, through a variety of mechanisms, including sequestration, efflux facilitation, and reduction of influx[19]. In any case, response variability to AgNPs among strains in nature or laboratory settings was expected[19]. Previous studies showed that fungi populations, including species like *Candida, Aspergillus, Cryptococcus*, and *Pneumocystis*, have different susceptibility rates to AgNP antifungal drugs[19,21]. It is important to prevent the development of resistance in the fungal population, whether by overdose or overuse of the compound, to avoid the selection of low susceptible strains.

This study also revealed that from concentrations of 25 mg L^-1^ to 50 mg L^-1^, more than 90% fungi growth inhibition was obtained. The concentrations of AgNPs that have been reported to inhibit *Fusarium* species vary across different studies. For instance, Singh et al. (2019) observed complete inhibition of *F. oxysporum* at 75 mg L^-1^ of AgNPs[22]. Similarly, Akpinar et al. (2021) suggested that AgNPs at 25 to 50 mg L^-1^ concentrations could effectively inhibit different *Fusarium oxysporum* f. sp. *radicis*-*lycopersici* strains[18]. Studies have shown that the size of AgNPs can influence their antifungal efficacy[18,23]. Additionally, the concentration of AgNPs used can impact their antifungal activity, with higher concentrations often leading to increased inhibition of fungal growth[12,24].

### *In vivo* interaction assay of AgNPs against *Fusarium*

The antifungal potential of the three types of AgNPs (Argovit-1220, Argovit-1221, and Argovit-C) was evaluated against fusarium wilt under greenhouse conditions using two AgNPs application: foliar and radicular, along with two methods of fusarium infection: root dipping and drenching. The percentage of infected areas revealed that AgNPs can control Foc race 1 in banana plants at a 100 mg L^-1^ concentration. Both ways of applications were effective in holding back the disease. Argovit-1220 had the highest antifungal potential followed by Argovit-C, which also showed a similar control. The least effective, according to the results, was the Argovit-1221.

The infection methods of Foc showed a difference between them, drenching infection achieved a greater infection area on the corm, even though several studies have reported better infection results with the root dipping method[25–27]. This difference could be explained because when using the root dipping infection method, many studies make 2 cm root cuts. Hence, the plant root wounds come in direct contact with the sporulated solution, making it easier for *Fusarium* spores to enter and colonize the root vascular tissue. However, in the experiment of the present work, we decided not to stress the plant in this way to not compromise the results of the experiment. It is important to note that the drenching method involved a longer contact time with the solution due to the use of polystyrene plates under the pots to retain the solution for a longer period.

Among the methods of application of the AgNPs, both the radicular application and foliar application had an effect as an antifungal agent. This means that it would be possible and easy to apply the AgNPs in the banana farms by the irrigation system and aerial spraying. In the case of radicular application, the AgNPs will be in direct contact with the pseudo-stem and the roots and will be absorbed by them easily. AgNPs will also be in contact with the soil, and since *Fusarium* is a soilborne fungus, the interaction between the AgNPs and the pathogen will be more direct than during the foliar one. As the foliar application also showed good results in controlling the disease, the foliar application is also considered a suitable method for spraying fungicide. It is important to emphasize that by applying directly to the leaf and not to the root, undesirable effects on soil micro-organisms community and the environment can be reduced. It is not known yet if there may be a significant diffusion and translocation of the AgNPs from the leaf to the root. Some studies report that AgNPs may act as a bio-stimulant that enhances the plant defence mechanisms against fungal infections[28–30].

The three types of AgNPs (Argovit-1220, Argovit-1221, and Argovit-C) were compared in this study under the same experimental variables; there was no statistical difference (p-value > 0.05) between the Argovit-1220 and the Argovit-C samples. Nevertheless, there was a statistical difference (p-value < 0.05) with the 1221 sample. The method of synthesis of AgNPs can also play an important role in their antifungal potential[31]. While the exact mechanisms are not fully understood, AgNPs are detrimental to fungal cell growth[32]. In general, the interaction of AgNPs with fungal cells has shown good controls for plant pathogenic fungi, suggesting their potential as an alternative to conventional fungicides[33]. We did not find any publication in the literature dedicated to the study *in vivo* assay of the use of AgNPs against *Fusarium* species. So, there is no way to compare our *in vivo* results with other literature data. Hence, according to our best knowledge, this work is the first *in vivo* assay of the use of AgNPs against *Fusarium oxysporum* f. sp. *cubense*.

Although AgNPs appear to be an excellent alternative for controlling this disease, some studies have raised concerns about the use of these compounds as future pesticides because of the environmental impact they may represent[34]. We suggest that more studies should be done to analyse the impact of AgNP application on the environment in agricultural settings.

## Conclusion

This study highlights the potential of AgNPs as effective fungicide agents against Foc race 1, both *in vitro* and *in vivo*. *In vitro* trial results indicated potent antifungal activity of AgNPs, with an average IC_50_ value of 4.33 ± 2.26 mg L^-1^ for Argovit-1220, 10.21 ± 8.44 mg L^-1^ for Argovit-1221, and 2.52 ± 1.13 mg L^-1^ for Argovit-C. Generally, it demonstrated high antifungal activity at low concentrations (12.5 mg L^-1^ – 25 mg L^-1^) against the different *Fusarium* strains. So, the Argovit-C AgNPs demonstrated *Fusarium* species growth inhibition 5 -10 times higher than earlier studied AgNP formulations. *In vivo* trial confirmed the potential of AgNPs to reduce the symptoms of Fusarium wilt disease in Gros Michel banana plants, controlling the infection in the corm of the plant and leading to improved plant health. At a concentration of 100 mg L^-1^ the radicular and the foliar applications of AgNPs showed an average inhibition of 61.83 ±18.5 and 68.83 ±12.53%, respectively.

It is important to emphasize that we did not find any research conducted on inhibiting *Fusarium oxysporum* f.sp. *cubense* race 1, whether *in vitro* or *in vivo*. So, according to our best knowledge, this is the first study demonstrating the systemic antifungal activity of AgNPs against Foc race 1 on banana plants. In addition, there is currently no conventional fungicide against Fusarium wilt, making these results relevant to the banana industry. This study underscores the importance of selecting appropriate application methods to maximize AgNP efficacy in controlling Foc infections. Further research is needed to optimize AgNP formulations and application protocols for field-scale implementation, and the impact of AgNPs on plant endophytes and the environment. This study contributes valuable insights into the development of innovative pesticide agents for managing fungal diseases in agriculture.

## Acknowledgments

We thank “FIASA - Fondo de Investigación para la Agrobiodiversidad, semillas y Agricultura Sustentable” for funding the research project (FIASA-CA-2023-008), ESPOL Polytechnic University (ESPOL), Vicerrectorado de Investigación de la Universidad de Cuenca (VIUC), The International Biotechnology Network supported by the “Consejo Nacional de Ciencia y Tecnología”, CONACYT, Mexico, and Russian Science Foundation and Tomsk Region Grant 22-13-20032, for funding this research.

## Notes

### Competing Interest Statement

The authors have declared no competing interest.

