## Supplement Table 1 for "Inhibition of *Fusarium oxysporum* growth in banana by silver nanoparticles: *in vitro* and *in vivo* assays": Supplemental Material-TableS1.docx

**S1 Table.** **Fungal growth inhibition percentages with different concentrations of AgNPs against three Fusarium oxysporum strains.**

| AgNPs concentrations [mg L^-1^] | Foc Strains | AgNPs | | |
| --- | --- | --- | --- | --- |
|  |  | **1220**  **Inhibition %** | **1221 Inhibition %** | **M Inhibition %** |
| 100 | Ec15  Ec19  Ec40  Ec35 | 100.00 ±0.00  100.00 ±0.00  100.00 ±0.00  100.00±0.14 | 96.24 ±4.69  99.18 ±1.10  98.14 ±1.63  84.64 ±8.42 | 99.87 ±0.00  99.65 ±0.00  99.52 ±0.00  84.26 ±5.89 |
| 50 | Ec15  Ec19  Ec40  Ec35 | 99.46 ±0.00  99.98 ±0.00  99.75 ±0.00  88.30 ±5.36 | 95.38 ±4.58  98.48 ±1.85  96.81 ±2.22  74.30 ±16.17 | 99.31 ±0.00  99.80 ±0.00  98.84 ±1.15  77.06 ±1.70 |
| 25 | Ec15  Ec19  Ec40  Ec35 | 98.43 ±0.58  99.12 ±0.58  98.15 ±1.15  80.21 ±12.14 | 86.64 ±3.25  95.26 ±4.50  87.10 ±9.64  60.31 ±14.99 | 95.56 ±2.31  98.03 ±2.31  93.75 ±2.08  66.62 ±3.43 |
| 12.5 | Ec15  Ec19  Ec40  Ec35 | 92.77 ±0.00  94.90 ±1.00  93.06 ±2.08  66.28 ±10.92 | 82.75 ±4.13  75.20 ±4.44  72.57 ±0.62  44.95 ±9.35 | 89.56 ±3.00  93.98 ±3.46  89.33 ±5.20  55.36 ±5.18 |
| 6.3 | Ec15  Ec19  Ec40  Ec35 | 82.12 ±3.06  82.07 ±4.16  78.91 ±8.14  48.91 ±8.10 | 59.66 ±11.59  51.24 ±4.29  57.28 ±10.08  36.84 ±11.88 | 73.56 ±4.04  85.79 ±3.00  78.24 ±5.51  48.27 ±8.32 |
| 3.1 | Ec15  Ec19  Ec40  Ec35 | 50.85 ±4.16  55.29 ±7.81  48.52 ±2.89  29.86 ±2.59 | 43.02 ±16.82  34.68 ±3.79  39.99 ±11.57  28.03 ±9.30 | 58.64 ±2.00  64.43 ±6.08  56.17 ±4.93  34.41 ±9.27 |
| 1.6 | Ec15  Ec19  Ec40  Ec35 | 32.68 ±1.73  32.02 ±3.06  29.04 ±2.52  19.87 ±5.65 | 30.11 ±19.78  17.42 ±1.88  25.46 ±9.90  22.79 ±9.08 | 40.60 ±1.15  48.45 ±6.08  38.02 ±1.73  25.37 ±7.89 |
| 0.8 | Ec15  Ec19  Ec40  Ec35 | 15.34 ±1.53  14.41 ±3.00  19.85 ±5.20  11.00 ±7.47 | 19.11 ±14.23  10.08 ±1.73  17.83 ±11.99  12.01 ±4.42 | 17.51 ±2.08  25.72 ±4.58  18.41 ±1.73  10.01 ±4.89 |
| 0 | Ec15  Ec19  Ec40  Ec35 | 0.00 ±0.02  0.00 ±0.04  0.00 ±0.01  0.00 ±0.01 | 0.00 ±0.00  0.00 ±0.01  0.00 ±0.02  0.00 ±0.01 | 0.00 ±0.34  0.00 ±0.30  0.00 ±0.34  0.00 ±0.01 |

Values are averages of 9 individual experiments. Data are given as mean values ± standard deviation.
